# starmapVR: immersive visualisation of single cell spatial omic data

**DOI:** 10.1101/2020.09.01.277079

**Authors:** Andrian Yang, Yu Yao, Xiunan Fang, Jianfu Li, Yongyan Xia, Crystal S. M. Kwok, Michelle C. K. Lo, Dickson M. D. Siu, Kevin K. Tsia, Joshua W. K. Ho

## Abstract

**Motivation:** Advances in high throughput single-cell and spatial omic technologies have enabled the profiling of molecular expression and phenotypic properties of hundreds of thousands of individual cells in the context of their two dimensional (2D) or three dimensional (3D) spatial endogenous arrangement. However, current visualisation techniques do not allow for effective display and exploration of the single cell data in their spatial context. With the widespread availability of low-cost virtual reality (VR) gadgets, such as Google Cardboard, we propose that an immersive visualisation strategy is useful.

**Results:** We present starmapVR, a light-weight, cross-platform, web-based tool for visualising single-cell and spatial omic data. starmapVR supports a number of interaction methods, such as keyboard, mouse, wireless controller and voice control. The tool visualises single cells in a 3D space and each cell can be represented by a star plot (for molecular expression, phenotypic properties) or image (for single cell imaging). For spatial transcriptomic data, the 2D single cell expression data can be visualised alongside the histological image in a 2.5D format. The application of starmapVR is demonstrated through a series of case studies. Its scalability has been carefully evaluated across different platforms.

**Availability and implementation:** starmapVR is freely accessible at https://holab-hku.github.io/starmapVR, with the corresponding source code available at https://github.com/holab-hku/starmapVR under the open source MIT license.

**Supplementary Information:** Supplementary data are available at *Bioinformatics* online.

## Introduction

Advances in single-cell RNA-seq technology (Ziegenhain *et al.*, 2017), flow cytometry (Saeys *et al.*, 2016), mass cytometry (Spitzer and Nolan, 2016), and quantitative phase imaging (Lee, Wang, *et al.*, 2019) has enabled the expression profiling of molecular expression and biophysical properties of hundreds of thousands of individual cells. Dimensionality reduction methods, such as Principal Component Analysis (PCA) or UMAP (Becht *et al.*, 2019), are typically used to reduce the large number of features into two or three dimensions to allow for visual exploration of the data. The clustering structure of single cells enable discovery of cell types and their potential relationships. Recently, attempts have been made to reconstruct 3D spatial organisation from scRNA-seq data (Ren *et al.*, 2020), which enables single cell data to be visualised in the context of their inferred 3D organisation. Moreover, spatial transcriptomic technology has enabled expression of multiple genes of thousands of cells in their 2D or 3D spatial context (Asp *et al.*, 2020). In many cases, it is a benefit to visualise the single cell omic data with the matching histological image, which provide important spatial context.

For both single cell and spatial omic data, a key analytical challenge is to be able to effectively explore global organisation of the cells, based on dimensionality-reduced expression space, and actual or inferred 2D or 3D organisation of cells. An effective visualisation should allow a viewer to not only assess the high-level spatial organisation of cells, but also to seamlessly ‘zoom’ into the data to discover fine local grouping of single cells, and to explore their molecular or phenotypic profiles, or in the context of additional spatial information such as the histological image in spatial transcriptomic. This type of visualisation is not easily achievable using standard visualisation tools. Inspired by the maturing field of immersive analytics (Marriott *et al.*, 2018), we believe adoption of interactive immersive visualisation techniques can improve the visualisation of single cell spatial data.

Here we present starmapVR as a light weight, cross-platform, web-enabled open source visualisation tool, that enables immersive visualisation of single-cell or spatial omic data for up to millions of cells. starmapVR can be launched on any device that has a web browser. When used on a mobile smartphone, starmapVR further utilises low-cost virtual reality (VR) headsets, such as Google Cardboard and Daydream, to enable widespread adoption of VR visualisation. We reason that an immersive visual experience will likely improve the intuitive navigation and exploration of hundreds of thousands of cells.

### Visual design, implementation, and applications

starmapVR breaks new ground on large scale single cell data and spatial transcriptomic data visualisation. It introduces a scalable visual design that combines the benefit of a three-dimensional (3D) scatter plot (Figure 1a) for exploring clustering or spatial structure and the benefit of star plots (Chambers *et al.*, 1983) (also known as radar chart) for multivariate visualisation of an individual cell (Figure 1b). starmapVR is built using web frameworks designed for creation of 3D and VR experiences, such as A-Frame and Three.js, which are cross-platform and can be adapted to computer screens and VR devices. Input data for starmap, in the form of 3D coordinates, cluster assignment, and value of up to 12 features per cell in a comma-separated values format, (and optionally one or more images for single cell imaging or spatial transcriptomic data), can be uploaded from a local file on your device or from third-party cloud storage, such as Google Drive, Microsoft OneDrive or iCloud Drive, depending on the support of the mobile smartphone’s operating system (see **Supplementary Materials** for detailed explanation of input format). starmapVR offers a number of options for navigation through the VR space. Users are able to use a wireless keyboard or wireless hand-held remote controller in order to scale, rotate and navigate within the VR space. starmapVR also offers a voice control feature that allows user to start and end custom animation for navigating the VR space.

**Figure 1.**
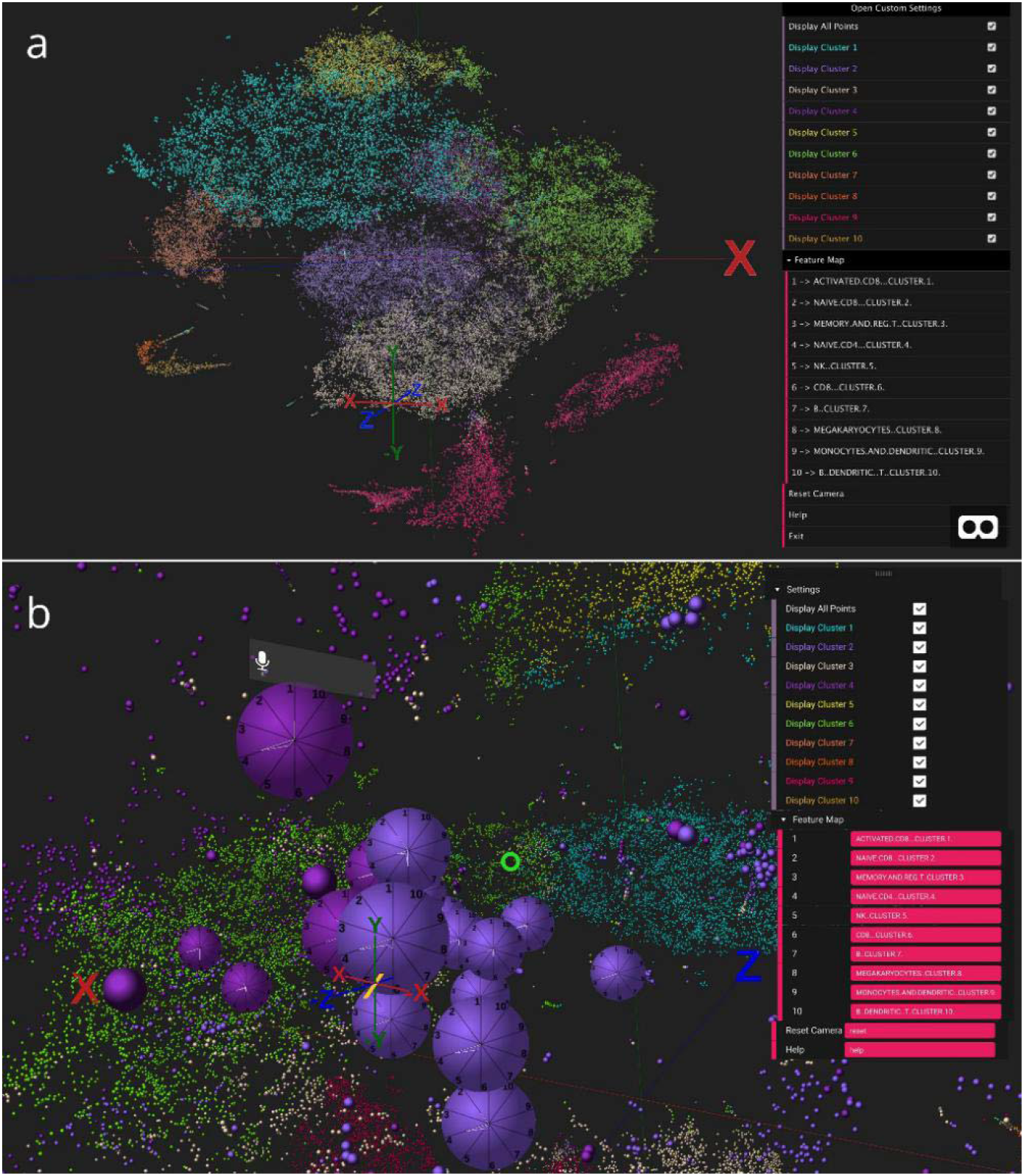
starmapVR visualisation of a single-cell RNA-seq data containing 68,000 peripheral blood mononuclear cells (**a**) Overview of the 3D scatter plot with visualised in non-VR mode (**b**) Star plot visualisation showing the relative expression of gene modules for each cell in VR mode.

For the star plot visualisation, starmapVR supports up to 12 radial coordinates which emanates from the centre to the circumference of the circle, with the coordinate indicating the expression level of a particular feature. The feature typically represents biological feature such a gene, in the case of RNA-seq data, a protein, in the case of flow cytometry data, or cellular phenotype, in the case of single cell imaging data. However, the feature can also represent can also be a loading value for a principal component, the mean expression of a group of genes or proteins, or some other aggregate representation of related features.

We argue that starmapVR can support visualise exploration of large single cell data in their inferred or actual spatial context. This can be highly valuable for bioinformaticians and other researchers. Since the use of starmapVR does not require expensive VR equipment, it may even be useful for visual communication of single cell data to the lay public.

To demonstrate starmapVR’s applicability, we have provided five example data sets on starmapVR website based on previously published single-cell RNA-seq data (Zheng *et al.*, 2017), flow cytometry data (The FlowCAP Consortium *et al.*, 2013), a reconstructed 3D spatial organisation inferred from single cell RNA-seq (Ren *et al.*, 2020), spatial transcriptomic data ( 10x Genomics) and single cell data with quantitative phase (QP) images (Lee, Lau, *et al.*, 2019). (see **Supplementary Discussion** for more details).

## Discussion

We released a preliminary version of starmapVR in 2018 (Yang *et al.*, 2018). Subsequently, more visualisation tools for single-cell data with VR technology have been developed. CellexalVR (Legetth *et al.*, 2019) is one of the VR visualisation platforms for single cell data. By placing all projected data calculated by different dimension reduction tools such as tSNE, diffusion maps and UMAP in VR, users can explore and compare different scRNA-seq analysing result. However, CellexalVR is designed to run on expensive VR hardware to operate, which limits its widespread usage. singlecellVR (Stein *et al.*, 2020) is developed to overcome the limitation of the old version of stramap by providing the package to convert outputs from commonly-used scRNA-seq analysis platforms such Scanpy and Seurat. Nonetheless, singelcellVR does not support spatial omic and single cell imaging data within their framework.

Tools such as the *spatial* package in Scanpy and ST-viewer (Fernández Navarro *et al.*, 2019) can be used to visualise spatial transcriptomic data. However, they only provide a very plain way to visualise these cells. starmapVR enables single cell multivariate data to be visualised alongside the spatial cues, such as a histological image that match the layout of the cells, in a two and a half dimensional (2.5D) visualisation strategy (Ware, 2001) (Supplementary Figure 6).

## Acknowledgement

We thank Dr Ken Yu and Dr Eleni Giannoulatou for providing critical comments on an earlier draft of the manuscript.

## References

Asp,M. et al. (2020) Spatially Resolved Transcriptomes—Next Generation Tools for Tissue Exploration. BioEssays, 1900221.

Becht,E. et al. (2019) Dimensionality reduction for visualizing single-cell data using UMAP. Nat. Biotechnol., 37, 38–44.

Chambers,J.M. et al. (1983) Graphical Methods for Data Analysis Wadsworth, Belmont, CA.

Fernández Navarro,J. et al. (2019) ST viewer: a tool for analysis and visualization of spatial transcriptomics datasets. Bioinformatics, 35, 1058–1060.

Lee,K.C.M., Lau,A.K.S., et al. (2019) Multi-ATOM: Ultrahigh-throughput single-cell quantitative phase imaging with subcellular resolution. J. Biophotonics, 12, e201800479.

Lee,K.C.M., Wang,M., et al. (2019) Quantitative Phase Imaging Flow Cytometry for Ultra-Large-Scale Single-Cell Biophysical Phenotyping. Cytometry A, 95, 510–520.

Legetth,O. et al. (2019) CellexalVR: A virtual reality platform for the visualisation and analysis of single-cell gene expression data. bioRxiv, 329102.

Marriott,K. et al. eds. (2018) Immersive Analytics Springer International Publishing.

Ren,X. et al. (2020) Reconstruction of cell spatial organization from single-cell RNA sequencing data based on ligand-receptor mediated self-assembly. Cell Res., 1–16.

Saeys,Y. et al. (2016) Computational flow cytometry: helping to make sense of high-dimensional immunology data. Nat. Rev. Immunol., 16, 449–462.

Spitzer,M.H. and Nolan,G.P. (2016) Mass Cytometry: Single Cells, Many Features. Cell, 165, 780–791.

Stein,D.F. et al. (2020) singlecellVR: interactive visualization of single-cell data in virtual reality. bioRxiv, 2020.07.30.229534.

The FlowCAP Consortium et al. (2013) Critical assessment of automated flow cytometry data analysis techniques. Nat. Methods, 10, 228–238.

V1_Human_Lymph_Node -Datasets -Spatial Gene Expression -Official 10x Genomics Support. Ware,C. (2001) Designing with a 2 1/2-D Attitude.

Zheng,G.X.Y. et al. (2017) Massively parallel digital transcriptional profiling of single cells. Nat. Commun., 8, 14049.

Ziegenhain,C. et al. (2017) Comparative Analysis of Single-Cell RNA Sequencing Methods. Mol. Cell, 65, 631–643.e4.

