## Supplementary Materials for "starmapVR: immersive visualisation of single cell spatial omic data"

**Supplementary Methods 1: Input data format for starmapVR**

starmapVR accepts as input a csv file or a zip-compressed csv file. The csv file need to contain a header row with the following column names - x, y, z and cluster - corresponding to the 3D coordinates of points and the cluster label assigned for each point (with outliers assigned the value of -1), see Supplementary Fig 1. In addition to the required columns, starmapVR also accepts extra columns (up to 12) corresponding to features which will be visualised in the star plot. The values for all columns must be of numeric types. For single cell imaging visualisation, two files are required: the same csv file as above and a compressed folder of single cell images the image file names using the numerical index of the cells. For spatial transcriptomic analysis, two files are required: a csv file and tissue histology image in which the coordinates in the csv file correspond with cell coordinates in the histology image. Examples of the input data used for case studies described below are available on starmapVR's GitHub repository.

| x | y | z | feature | feature | feature | . . . | feature | feature | feature | feature | cluster |
|---|---|---|---|---|---|---|---|---|---|---|---|
| | | | | | | . . . | | | | | |
| | | | | | | . . . | | | | | |
| | | | | | | . . . | | | | | |
| | | | | | | . . . | | | | | |
| | | | | | | . . . | | | | | |

**Supplementary Figure 1.** Sample input file format, where there can be up to 12 features, "feature" can be changed into customized column names.

**Supplementary Methods 2: Compatibility with Scanpy and Seurat**

Scanpy and Seurat are among the most widely-used single-cell analysis platforms. We provide simple functions that convert single cell RNA-seq objects in Scanpy and Seurat to the required input format for starmapVR. Users can choose any coordinate system for the single cells, such as UMAP, t-SNE or PCA coordinates. Features for star plots can be based on any user-specified values, such as expression of a handful of selected genes, selected principal components, or some composite scores. The single cells in starmapVR can be coloured based on cell cluster labels or expression value for a selected gene. An example of using starmapVR to visualise and analysis the PBMC data set of 3,962 cells provided as part of the Scanpy and Seurat tutorial is shown in Supplementary Fig 2. We also provide the example that can transform the spatial omic data object in Scanpy to the desired input for starmapVR, more detail can be found in Case Study 4.

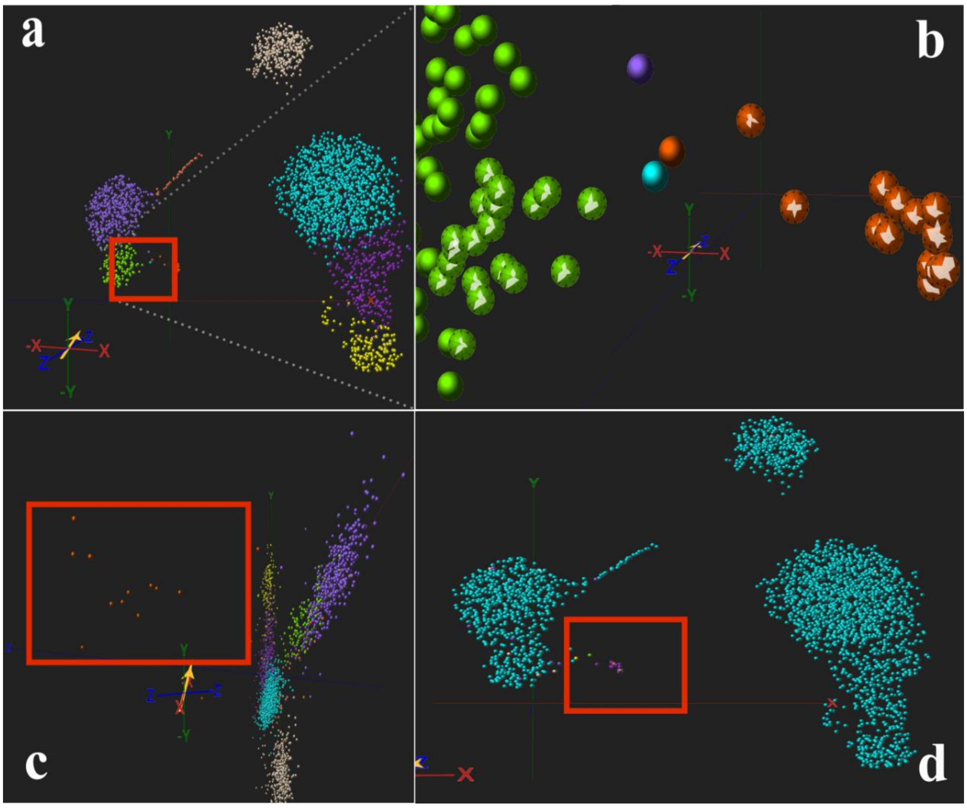

**Supplementary Figure 2.** starmapVR visualisation of the PBMC data set used in Scanpy & Seurat tutorial. (a) Overview of the 2D scatter plot, cluster 7 of interest is highlighted. (b) Star plots of the area of interest for cluster 7 (orange colour) and cluster 5 (green colour), cluster 7 has a significant higher value of PC3 compared to cluster 5. (c) 3D visualisation using PC1, PC2 and PC3 as coordinates, most of the cells fall in the x-y plane (PC1 and PC2), while cells from cluster 7 spread with z-axis (PC3) (d) 2D scatter plot of PBMC data set coloured by *PF4* gene (the highest weight gene contributed to PC3), cells with blue colour are cells with *PF4* expression level < 1.

**Supplementary Methods 3: Interaction in the virtual reality space in starmapVR**

starmapVR supports a number of input methods for interacting with the visualisation - keyboard, remote control and voice control. Note that voice control is available only in Google Chrome (desktop and mobile) as voice control utilises the SpeechRecognition API which is currently only supported by Google Chrome web browser. A summary of the control scheme for keyboard and voice control is included in Supplementary Table 1, while the control scheme for remote control is shown in Supplementary Fig 3.

**Supplementary Table 1.** Keyboard control scheme and voice control commands for starmapVR.

| Command | Keyboard Control | Voice Command |
|---|---|---|
| forward | w | forward |
| backward | s | backward |
| left | a | left |
| right | d | right |
| zoom in | q | in |
| zoom out | e | out |
| rotate Y-axis clockwise | left arrow | N\A |
| rotate Y-axis anti-clockwise | right arrow | rotate |
| rotate X-axis clockwise | up arrow | N\A |
| rotate X-axis anti-clockwise | down arrow | N\A |
| click on toolbox (VR mode) | N\A | select |
| reset toolbox (VR mode) | N\A | reset |
| reset view (VR mode) | N\A | init |

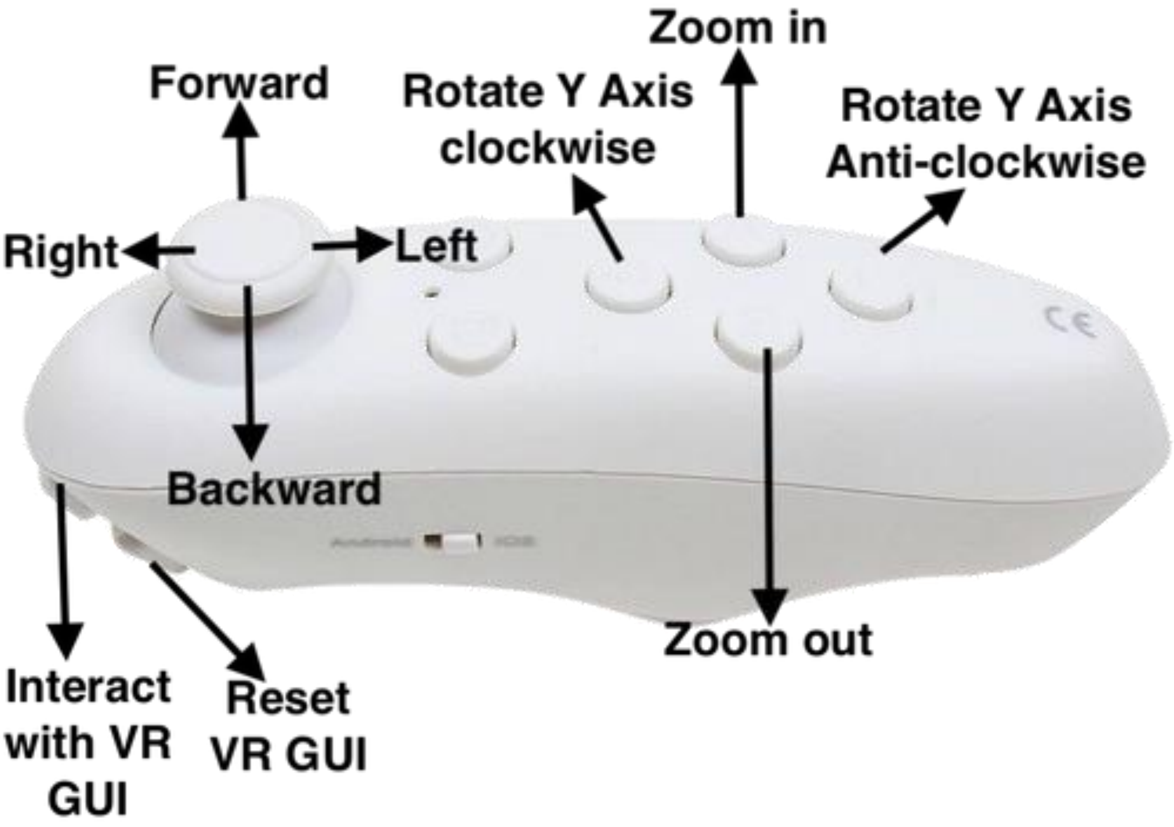


**Supplementary Figure 3.** Control mapping for remote control in starmapVR.

**Supplementary Methods 4: Performance evaluation**

We evaluated the performance of starmapVR on two key metrics – data loading time and rendering performance. The data set utilised for the performance testing is a synthetic data set with varying number of points – ranging from 200,000 up to 1,500,000 – generated using Python programming language. The web browser utilised for testing in both mobile and desktop devices is Google Chrome browser.

In the loading time performance test, we evaluated the time taken by starmapVR to load a file from the local device, process the data and render the visualisation. We selected this metric as the size of single-cell data set is in the order of hundreds of thousands and thus it is important that starmapVR is able to quickly load and visualise the data. We utilised files from local device to measure only the performance of starmapVR in loading and processing the data. We defined the loading time as the time difference between the start of file upload and the end of the scene render. We chose to evaluate this metric as the number of points in single-cell data set will be in the order of hundreds of thousands and thus it is important to ensure that starmapVR is able to quickly load the data.

The result of the loading time performance test in both mobile and desktop devices shows that newer devices are able to load data faster than older devices, due to the increase in processing power (both CPU and GPU) on newer devices (Supplementary Fig 4 and 5). The result of the performance test also shows that desktop devices are able to load the data faster than mobile devices, with all desktop devices able to load the largest data set containing 1,500,000 points in ≤ 6 seconds.

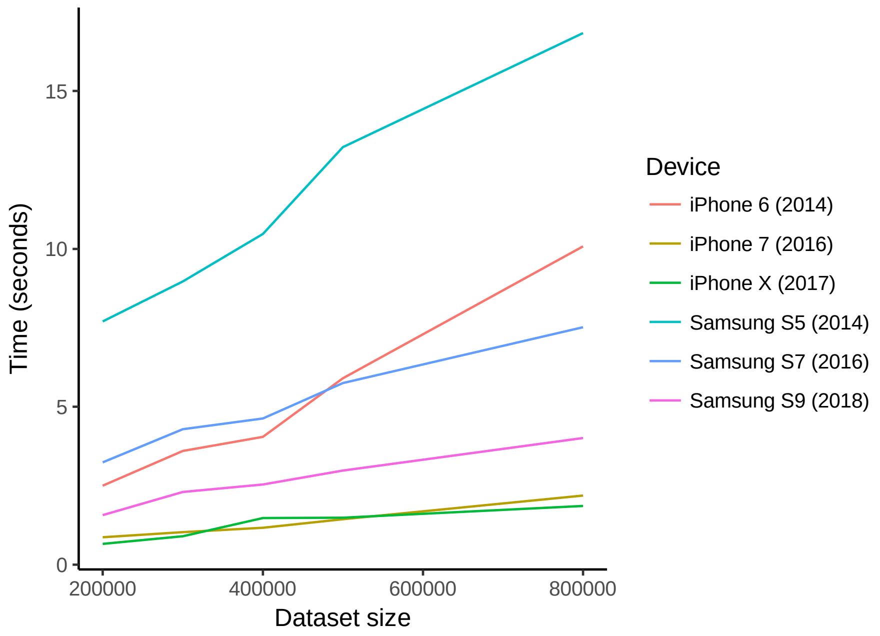


**Supplementary Figure 4.** Loading and processing time of varying data set sizes on mobile devices.

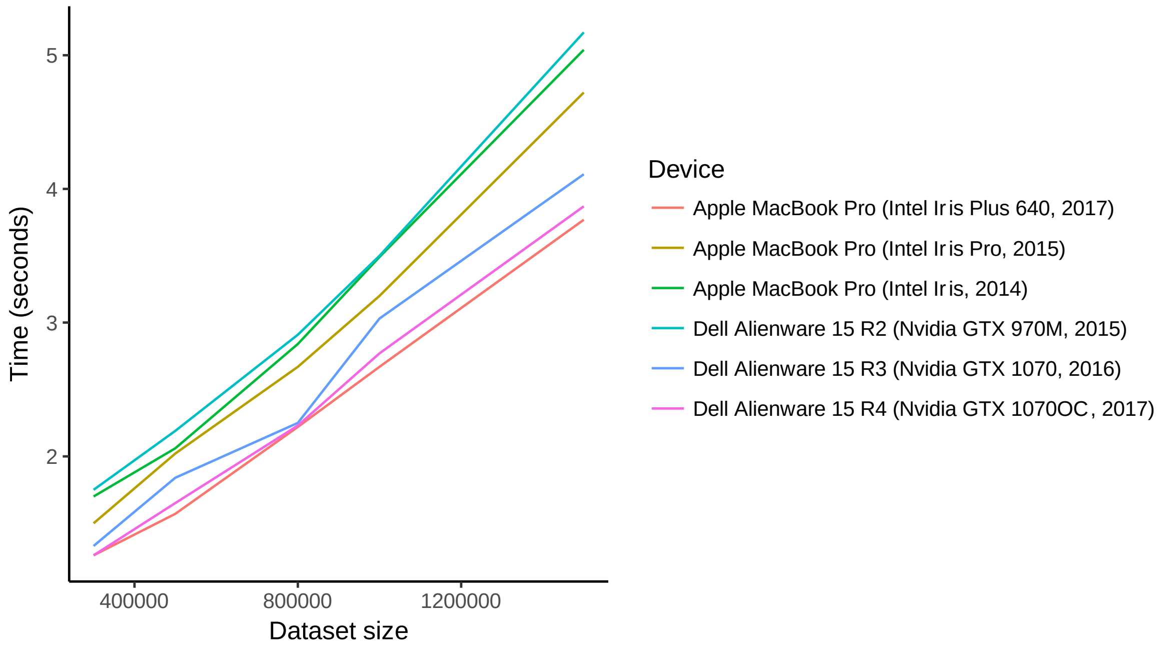


**Supplementary Figure 5.** Loading and processing time of varying data set sizes on desktop devices.

For the rendering performance test, we evaluated the ability of the device at maintaining a stable visualisation at 30 frame per seconds (FPS) using the statistics component of A-Frame.js to monitor the framerate rendered on screen. The result of the rendering performance test shows a similar trend as the previous test, with newer devices able to successfully achieve a framerate of 30 FPS due to better processing power on newer devices (Supplementary Tables 2 and 3). Furthermore, the result of the test shows that desktop devices are able to render larger data set at a stable 30 FPS compared to mobile devices, due to the greater processing capability of desktop GPUs compared to mobile GPUs as starmapVR utilises GPU acceleration.

**Supplementary Table 2.** Rendering performance test on mobile devices with varying data set sizes. Checkmarks indicates that the mobile device is able to render the data set with a minimum of 30 frames per seconds.

| Mobile device (name, year) | Data set size | | | | |
|---|---|---|---|---|---|
| | 200,000 | 300,000 | 400,000 | 500,000 | 800,000 |
| iPhone 6 (2014) | ✓ | ✓ | ✓ | | |
| iPhone 7 (2016) | ✓ | ✓ | ✓ | ✓ | |
| iPhone X (2017) | ✓ | ✓ | ✓ | ✓ | ✓ |
| Samsung S5 (2014) | ✓ | ✓ | ✓ | ✓ | |
| Samsung S7 (2016) | ✓ | ✓ | ✓ | ✓ | ✓ |
| Samsung S9 (2018) | ✓ | ✓ | ✓ | ✓ | ✓ |

**Supplementary Table 3.** Rendering performance test on mobile devices with varying data set sizes. Checkmarks indicates that the mobile device is able to render the data set with a minimum of 30 frames per seconds.

| Desktop device (name, GPU, year) | Data set size | | | | |
|---|---|---|---|---|---|
| | 300,000 | 500,000 | 800,000 | 1,000,000 | 1,500,000 |
| Apple MacBook Pro (Intel Iris, 2014) | ✓ | ✓ | ✓ | | |
| Apple MacBook Pro (Intel Iris Pro, 2015) | ✓ | ✓ | ✓ | | |
| Apple MacBook Pro (Intel Iris Plus 640, 2017) | ✓ | ✓ | ✓ | ✓ | ✓ |
| Dell Alienware 15 R2 (Nvidia GTX 970M, 2015) | ✓ | ✓ | ✓ | ✓ | |
| Dell Alienware 15 R3 (Nvidia GTX 1070, 2016) | ✓ | ✓ | ✓ | ✓ | ✓ |
| Dell Alienware 15 R4 (Nvidia GTX 1070OC, 2017) | ✓ | ✓ | ✓ | ✓ | ✓ |

### Supplementary Discussion

#### Case study 1: 3D t-SNE visualisation of single-cell RNA-seq data

We utilised the 68,000 peripheral blood mononuclear (PBMC) single-cell RNA-seq data from Zheng *et al.*[1] to showcase the ability of starmapVR to visualise the entire data set using a 3D t-SNE projection, grouped into 10 cluster, along with the expression of gene modules in each single cell as a star plot. We obtained the gene expression profile for the PBMCs and the source code utilised previously for analysis of the PBMC data from 10XGenomics GitHub repository (https://github.com/10XGenomics/single-cell-3prime-paper). In short, gene profiles were normalised according to the barcode (UMI) counts, followed by principal component analysis on the top 1000 most variable genes. The PCA values were then used to perform t-SNE to produce 3D coordinates and *k*-means clustering to obtain 10 clusters. For gene modules expression of individual clusters, we calculated the $\log_2$-transformed mean normalised expression of the genes specific to each cluster for all 10 clusters.

#### Case study 2: Single cell flow cytometry data

We used a flow cytometry data set from cell populations of HIV-exposed uninfected and un-exposed infants generated by Aghaeepour *et al*[2]. We combined the measurements from 10 selected samples to obtain a combined 4,000,000 cells measurements of 8 biomarkers. The data were then adjusted to truncate negative values to zeroes and hyperlog-transformed to obtain a normalised measurement. Clustering was then performed to group the cells into 3 clusters and principal component analysis were used to reduce the dimension of the data for visualisation. Due to the large number of cells within the data set, stratified sampling was used to down sample the data to 500,000 cells.

**Case study 3: Visualisation of inferred 3D spatial organisation of single cells based on ligand-receptor self-assembly (CSOmap)**

One major area of research in single cell analysis is the inference of three-dimensional spatial arrange of single cells, the so-called pseudo-space analysis, using single cell omic data. Ren et al. [3] recently developed a new computational method called CSOmap to infer cellular spatial organisation based on ligand-receptor interactions using scRNA-seq data. Considering the known benefit of VR over traditional 3D visualisation in exploration of spatial data, we believe starmapVR will be useful in visualising the CSOmap inferred 3D spatial data. As a demonstration, we applied starmapVR to visualise the CSOmap inferred 3D coordinates of 4,513 single cells in a melanoma data set from Tirosh *et al*[4]. We specified the first ten principal components to be visualised on each cell's star plot, based on the 23,686 genes in the data set. As show in Supplementary Fig. 6, starmapVR provides a straight-forward means to compare the expression levels (as represented by the star plots of the first 10 PCs) of cells that are determined to be spatially nearby in the inferred pseudospace.

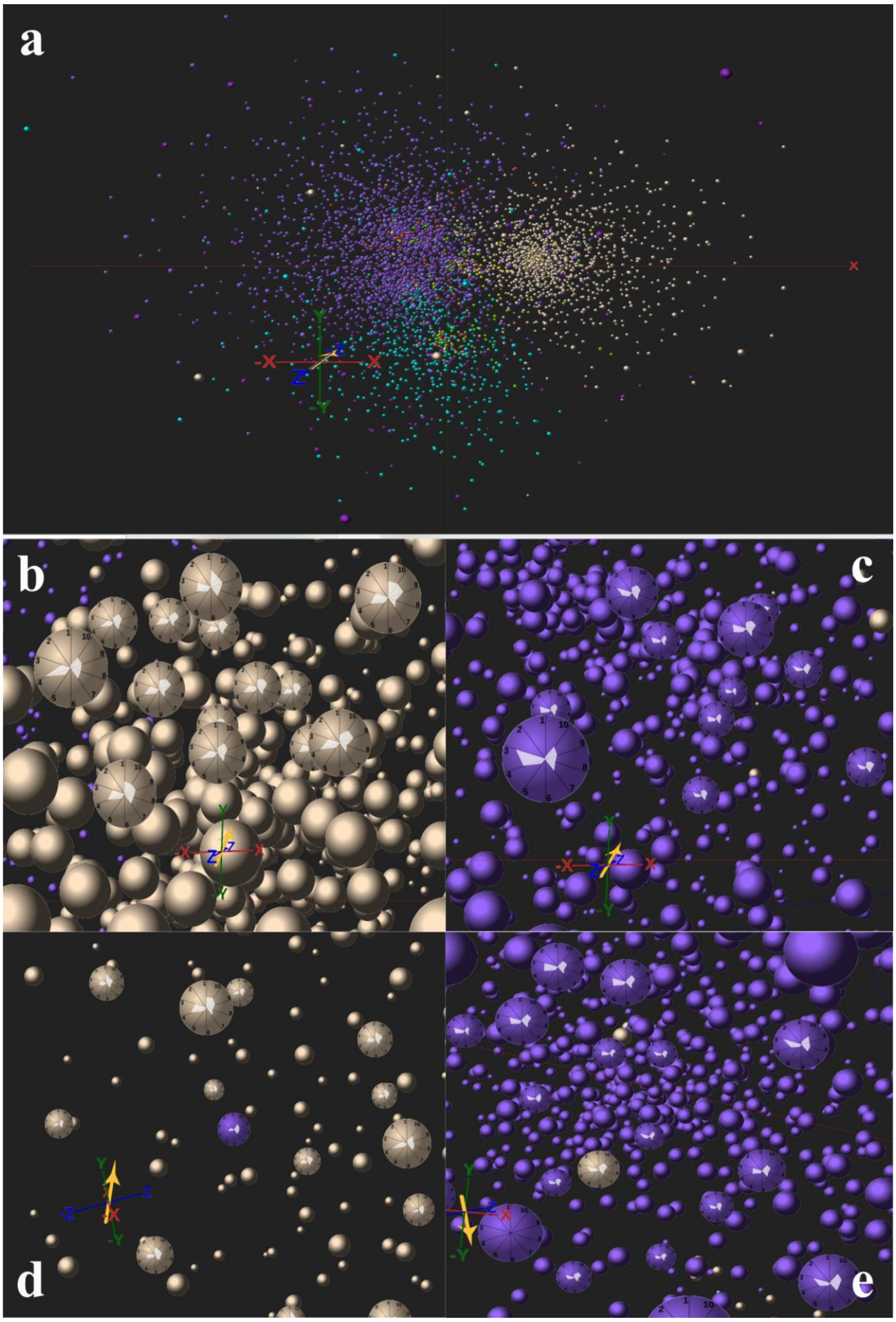


**Supplementary Figure 6.** starmapVR visualization of melanoma data set used in CSOmap. (a) Overview of starmapVR visualization of the melanoma data set. (b) Star plots of malignant cells (beige colour) in a homogeneous niche, (c) Star plots of T cells (purple colour) in a homogenous niche. (d) T cells in malignant cell niche. (e) Malignant cells in T cell niche.

At a global level, malignant cells (beige colour) and T cells (purple colour) occupy distinct regions in the 3D pseudospace (Supplementary Fig 6a). When we closely examined and compared the star map of the malignant cells and T cells using starmapVR, we noticed that the main difference between these two cell types are in the value of PC3 (higher in T cells and lower in malignant cells; Supplementary Fig. 6b and 6c). Interestingly, by focusing on some of the individual cells that are surrounded by other cell types, we observed the annotated T cells lost the characteristic high PC3 value (Supplementary Fig. 6d), and the malignant cells lost its characteristic low PC3 value (Supplementary Fig.  6e). These observations suggest that perhaps some of the cell type annotation is not correct, or that PC3 can be affected by the cellular environment.  By analysing the key genes that contribute to PC3, the gene with the highest weight is *B2M*, beta-2 microglobulin, which has a potential role as a convenient and non-invasive prognostic indicator in malignant lymphomas[5]. This case study highlight how starmapVR's ability to interactively interrogate both the global spatial organisation of cells and the expression patterns of individual cells can reveal interesting biological observations.

**Case study 4: 2.5D visualisation of spatial transcriptomic data**

Spatial transcriptomics enables the study of transcriptional activity of single cells in the context of spatial organisation of these cells in the endogenous tissue environment. By adding spatial information to scRNA-seq data, spatial transcriptomics enables researchers to understand cell-to-cell interactions in their native environment. Thus new tools which are capable of better visualizing data with the extra spatial dimension are necessary to unveil the potential information contained in this novel type of data. We demonstrate how to visualise a spatial transcriptomic object in Scanpy with starmapVR, using a public available Visium spatial transcriptomics data set with 4,039 cells of the human lymph node [6]. Cells are overlaid on top of the hematoxylin and eosin (H&E) stained histology image provided (Supplementary Fig. 7a). Cells are visualised in spatial coordinates and coloured by Leiden clustering result based on gene expression matrix (Supplementary Fig.7). By comparing cells clusters calculated by gene expression matrix and original tissue image, the organisation of cell types inferred by gene expression is consistent with the cellular arrangement identified by the histological image. For example, cells in cluster 7 (orange colour) are overlap with the light purple regions in the H&E image (Supplementary Fig. 7c) and surrounded by cells belonging to Cluster 1(blue colour) (Supplementary Fig. 7d). By comparing star plots of cluster 7 and 1 with other clusters, we noticed that cells from these two groups have a higher PC2 value, and the gene with the highest weight is *CR2*, Complement receptor type 2. Cells coloured by *CR2* gene expression level 5 and 3 show a very similar spatial distribution with cluster 7 and 5. By visualising clustered samples in spatial dimensions, we can gain insights into tissue organisation and potential inter-cellular communication.

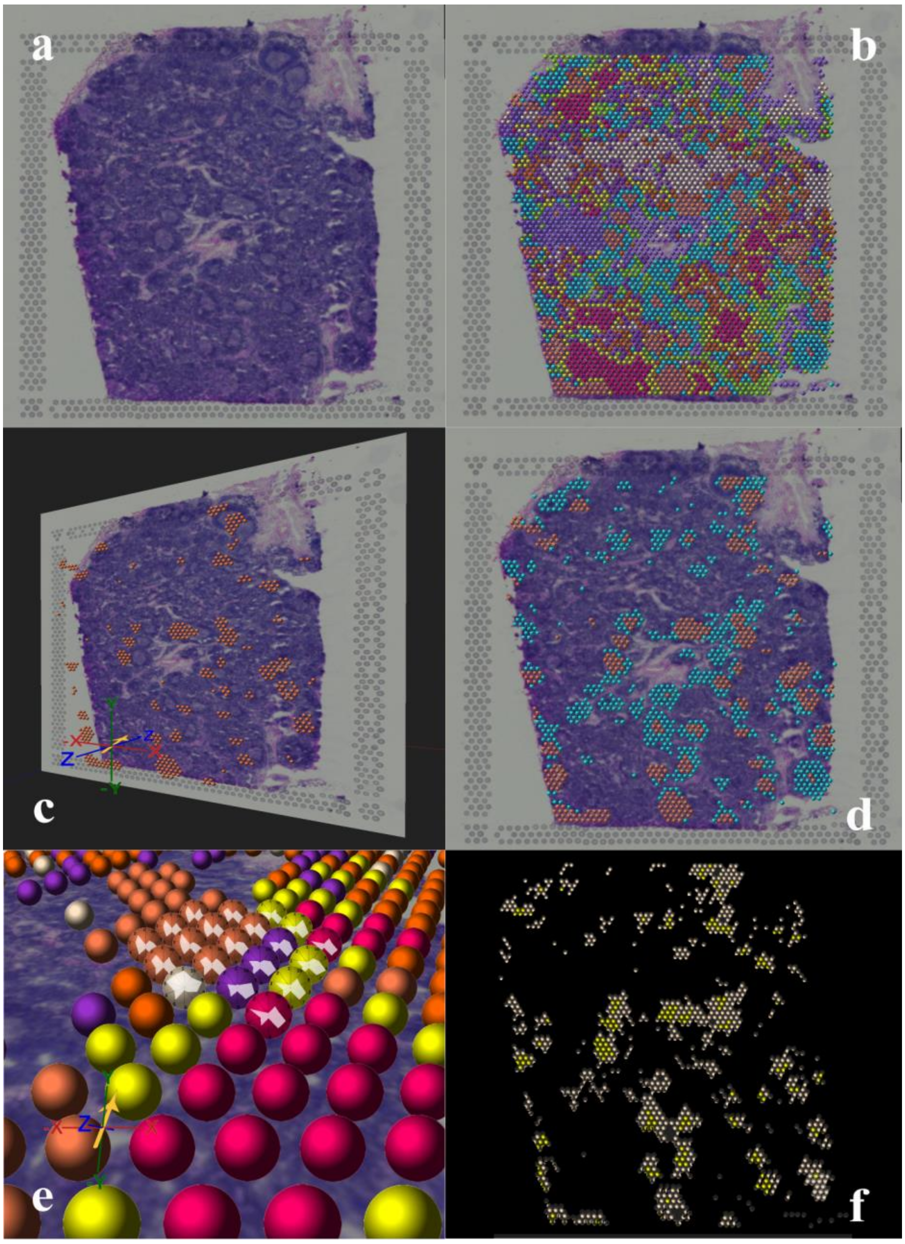


**Supplementary Figure 7.** Spatial transcriptomic data analysis using starmapVR. (a) Hematoxylin and eosin (H&E) stain image the human lymph node. (b) Spatial data coloured according cell types identified based on gene expression data. (c) Cells in cluster 7 with orange colour correlated with H&E image. (d) Cells from cluster 7 (orange colour) surrounded by cells belonging to Cluster 1(blue colour). (e) Star plots of cells from cluster 7 and neighboring cells. (f) Spatial data colored according to the *CR2* gene expression level.

**Case study 5: Visual analysis of biophysical phenotype profiling data by single cell imaging**

Cellular biophysical properties are the effective label-free phenotypes indicative of differences in cell types, states, and functions. starmapVR enables visualisation of single-cell quantitative phase (QP) images (captured by an ultrahigh-throughput imaging technique called Multi-ATOM[7]), in the context of their biophysical phenotypic profiles[7]. The QP image represents the spatial distribution of dry mass density within the cells, from which a large number of biophysical phenotypes can be derived. We utilise a Multi-ATOM data set which contains 105,000 cells from seven lung cancer cell lines (15,000 cells per cell line): i.e., three adenocarcinoma cell lines (H358 [EGFR WT], HCC827 [EGFR exon 19 del] and H1975 [L858R and T790M]) two squamous cell carcinoma cell lines (H520 and H2170), two small cell lung cancer cell lines (H526 and H69). For star plot visualisation, we highlight ten biophysical phenotypes of interest: volume, dry mass, peak of phase, variance of phase, phase centroid displacement, QP entropy mean, QP entropy variance, QP entropy skewness, QP entropy kurtosis and QP entropy radial distribution. The spatial coordinates in starmapVR are the first three PCs from PCA based on the 84 extracted biophysical features from Multi-ATOM. In Supplementary Fig. 8, the star plots are replaced by the single cell QP images (colour coded with the quantitative phase values[7]).

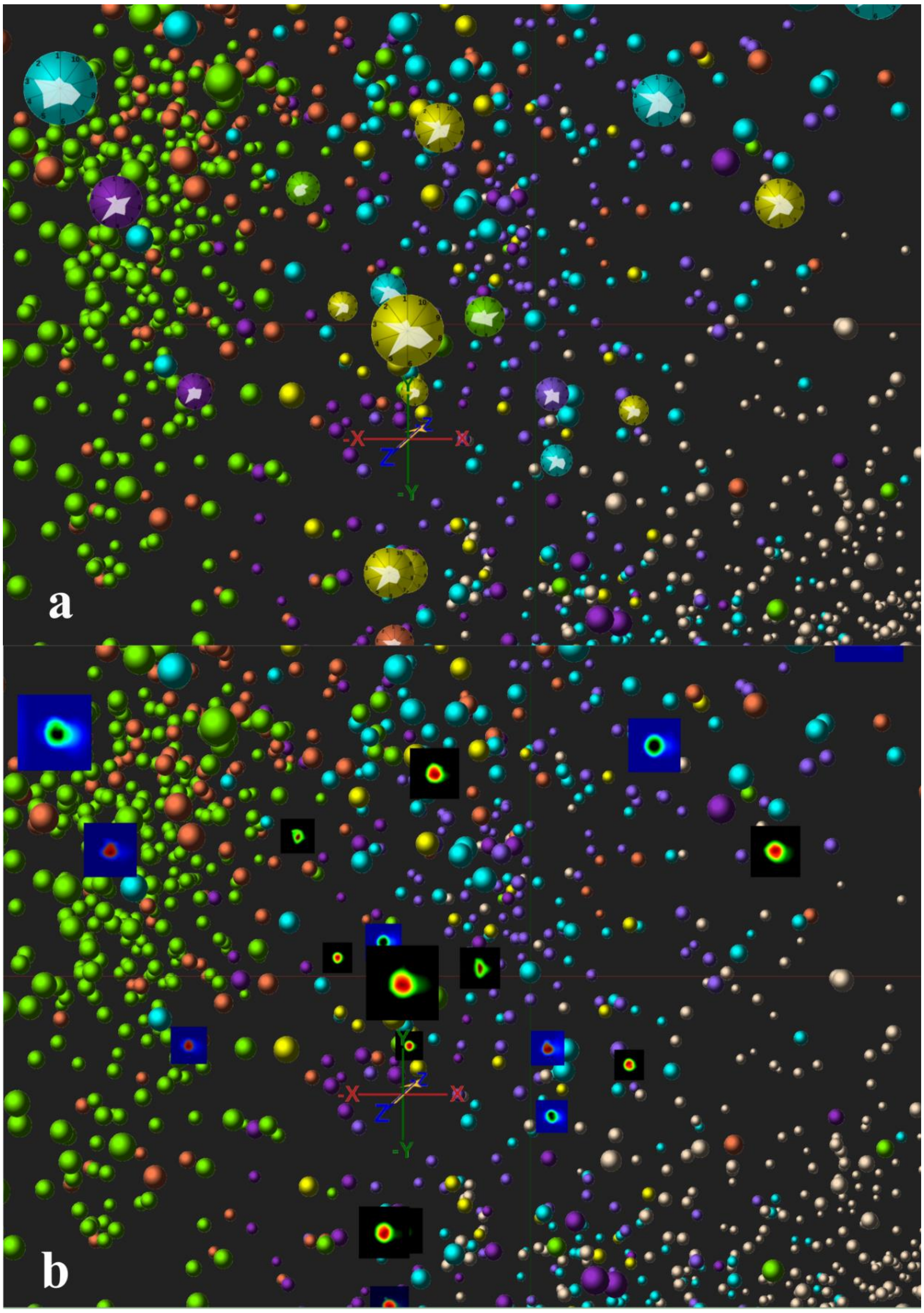


**Supplementary Figure 8.** Cells with real images visualised in starmapVR. (a) Normal star plot visualisation. (b) Real cell image in replace of star plot.

### Comparison with other relevant visualisation tools

**Supplementary Table 3.** Comparison with other relevant visualisation tools

| | **starmapVR** | **CellexalVR** | **SinglecellVR** |
|---|---|---|---|
| Web browser support | Yes | No | Yes |
| Smartphone support | Yes | No | Yes |
| Compatibility with Scanpy and Seurat | Yes | No | Yes |
| Comparison of multiple data sets | No | Yes | No |
| Spatial transcriptomic data visualisation | Yes | No | No |
| Image cytometry data visualisation | Yes | No | No |

**References**


1. Zheng, G. X. Y. *et al.* Massively parallel digital transcriptional profiling of single cells. *Nat. Commun.* **8**, 14049 (2017).

2. The FlowCAP Consortium *et al.* Critical assessment of automated flow cytometry data analysis techniques. *Nat. Methods* **10**, 228–238 (2013).

3. Ren, X. *et al.* Reconstruction of cell spatial organization from single-cell RNA sequencing data based on ligand-receptor mediated self-assembly. *Cell Res.* 1–16 (2020) doi:10.1038/s41422-020-0353-2.

4. Tirosh, I. *et al.* Dissecting the multicellular ecosystem of metastatic melanoma by single-cell RNA-seq. *Science* **352**, 189–196 (2016).

5. Yoo, C., Yoon, D. H. & Suh, C. Serum beta-2 microglobulin in malignant lymphomas: an old but powerful prognostic factor. *Blood Res.* **49**, 148–153 (2014).

6. V1_Human_Lymph_Node -Datasets -Spatial Gene Expression -Official 10x Genomics Support. https://support.10xgenomics.com/spatial-gene-expression/datasets/1.0.0/V1_Human_Lymph_Node.

7. Lee, K. C. M. *et al.* Multi-ATOM: Ultrahigh-throughput single-cell quantitative phase imaging with subcellular resolution. *J. Biophotonics* **12**, e201800479 (2019).
